# Body-map proto-organization in newborn macaques

**DOI:** 10.1101/565390

**Authors:** Michael J. Arcaro, Peter F. Schade, Margaret S. Livingstone

## Abstract

Topographic sensory maps are a prominent feature of the adult primate brain. Here, we asked whether topographic representations of the environment are fundamental to early development. Using fMRI, we find that the newborn somato-motor system, spanning frontoparietal cortex and subcortex, comprises multiple topographic body representations. The organization of these large-scale body maps was indistinguishable from those in adults and already exhibited features stereotypical of adult maps. Finer-scale differentiation of individual fingers increased over the first two years, suggesting that topographic representations are refined during early development. Last, we found that somato-motor representations were unchanged in two visually impaired monkeys who relied entirely on touch for interacting with their environment, demonstrating that massive shifts in early sensory experience in an otherwise anatomically intact brain are not sufficient for driving cross-modal plasticity. We propose that a topographic scaffolding is present at birth that both directs and constrains experience-driven modifications throughout sensory systems.

## INTRODUCTION

Smooth and continuous representations of the sensory periphery, i.e. topographic maps, are a fundamental feature of information processing in the adult primate brain. Topographic maps cover most of the cortical surface in primates. The adult primate somatosensory and motor systems comprise several areas spanning parietal and frontal cortices, as well as subcortex. Each area contains a topographic map of the body ^1-4^. This large-scale organization, as well as the anatomical and functional properties of individual areas, are similar across individuals, suggesting a common program for their development. However, the mechanisms guiding such organization remain unresolved. Prior microelectrode recordings in newborn macaques found neurons in primary somatosensory cortex (3a/b) to be unresponsive to tactile stimulation and immature in infant marmosets ^5^. Further, area 1, which processes information from area 3a/b ^6^, was less responsive in newborns than in older monkeys, all of which suggests that the primate somato-motor system is functionally immature at birth. Beyond these early somatosensory areas, the functional organization of the somato-motor hierarchy at birth has yet to be explored.

Here, we ask whether somatotopic representations of the body are present at birth, and to what extent does the organization of the newborn somato-motor system resemble that found in adults. Using fMRI, we show that frontal and parietal cortices, as well as subcortical structures, respond to stimulation of the face, hand, and foot in newborn monkeys. These representations are organized into multiple large-scale topographic maps of the body. The organization of these maps was indistinguishable from maps found in older monkeys and contained stereotypical features such as the overrepresentation of the face and hands in primary somato-motor cortex. Coarse representations of the fingers were present at birth and were refined over the first two years of postnatal development. We found a comparable organization in two monkeys that were raised from birth under visual form deprivation. These monkeys behaviorally rely on somatosensation for navigation and interacting with their environment, in contrast to typically reared monkeys that rely primarily on vision. The lack of large-scale reorganization suggests that extreme shifts in early postnatal sensory experience are not sufficient for cross-modal remapping of somatotopic representations. Together, these results show that areal differentiation and the topographic layout of the somato-motor system is present at birth, and likely serves as the scaffold for subsequent postnatal development.

## RESULTS

### Cortical responses to tactile stimulation in newborns and juveniles

We performed somatotopic (tactile body) mapping in nine monkeys ranging in age from 11 to 669 days. Each monkey exhibited robust cortical responses to tactile stimulation of the contralateral face, hand, and foot (*p*<0.0001 uncorrected, Figure 1; Supplementary Figure 1). Within the central sulcus, activity varied along the dorsal / ventral axis with face responses localized to ventral-most regions, foot and lower back responses to dorsal-most regions, and hand responses falling in between. These activity patterns to contralateral stimulation were consistent across monkeys (Figure 1 top; Supplementary Figure 1). For a subset of monkeys, the lower back was also stimulated (Supplementary Figure 1). Evoked activity from lower back stimulation overlapped activity from stimulation of the foot, but was also shifted ventral within the central sulcus above the hand representation. The relative location of these responses is consistent with prior electrophysiology, fMRI, and stimulation studies in macaques demonstrating spatially distinct representations of the face, hand, and foot within primary somatosensory ^7-12^ and motor cortex ^2, 13, 14^. Even in the youngest monkey tested (M1, 11 days), tactile stimulation evoked spatially specific activity within the central sulcus (Supplementary Figure 1), demonstrating that the functional responses of primary somato-motor cortex to tactile stimulation of the contralateral body shortly after birth are already spatially specific along the cortical surface.

**Figure 1.**
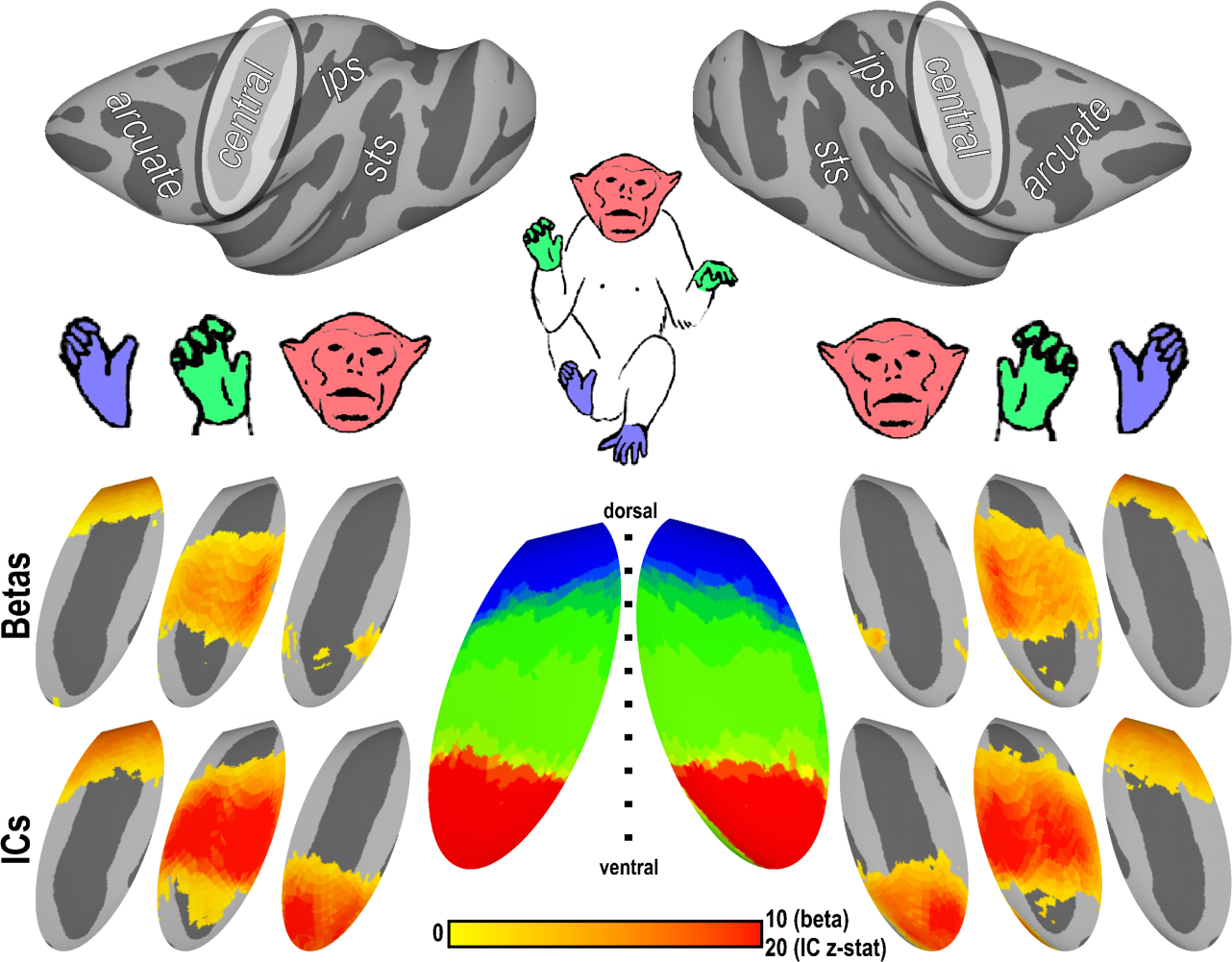
Somatotopic organization of primate central sulcus in juveniles. Spatially-specific responses within the central sulcus were observed to stimulation of the contralateral foot, hand, and face from both GLM (beta values) and independent component (IC) analyses. Group average (n = 9 in right hemisphere and n = 7 in the left hemisphere) data show vertices that were significantly active (p < 0.0001, uncorrected) in at least 4 subjects from a conjunction analysis (Supplementary Figure 1). Combined, these activity maps formed an inverted topographic gradient of body representations (center) with the face (red) represented ventrally, feet (blue) represented dorsally, and a large hand representation (green) in between. Monkey illustrations adapted from ^15^.

During scanning the monkeys were fully alert. Though each monkey was head restrained, their limbs were unrestrained and could move freely within the primate chair. It is likely that self-generated movement added noise in our regression analysis for estimates of body-part evoked activity because predictors were labeled based solely on when we stimulated their body parts. To better detect brain areas that reflect all activity related to stimulation of each body part, we conducted an independent component analysis (ICA) on the data (Methods: Independent Component Analysis (ICA)). This data-driven approach can identify significant structure in the data that is not isolated solely to the condition blocks. e.g., a component may capture hand movement-related activity both during and between hand stimulation blocks. Spatial activity maps were then reconstructed from each independent component to visualize the regions of the brain whose signals most contributed to each component. Components corresponding to the foot, hand, and face regions within the central sulcus were identified in each monkey (Figure 1, bottom; Supplementary Figure 2). There was a strong correspondence between the anatomical location of activity (beta) maps from the regression analysis and IC maps for both feet and hands. Across monkeys, hand and most leg IC maps were unilateral and corresponded to beta maps of contralateral stimulation. In each monkey, the face IC map was bilateral and symmetric. The face IC map included the face beta maps for both hemispheres, but was more extensive, encompassing the entire ventral third of the central sulcus. We attribute this difference to the fact that the monkey’s head and face were continuously stimulated throughout each scan, including baseline periods, from the helmet restraint and receipt of periodic juice reward. While the group average beta and IC maps were similar for each body part, the IC maps were more consistent and anatomically focal across individuals, indicating that this analysis better captured body-part specific activity throughout the experiment.

### Somatotopic organization of primary somato-motor cortex in newborns

A topographic map of the body was identified in newborn monkeys as early as 11 days of age. To identify topographic body maps, the weights of face, hand, and foot ICs were used to derive the region of the body each voxel most strongly represented (Methods: Body Map Analysis). Consistent with the activity maps for individual body parts, a topographic gradient of body part representations was identified within the central sulcus (Figure 1, middle). A large representation of the face (red) was identified in the ventral third of the central sulcus from the body map analysis. This representation progressed to a hand representation (green) in the middle of the central sulcus and then to a representation of the foot dorsally (blue). These data demonstrate the presence of a topographic map of the body within primary somatosensory and motor cortex of newborn monkeys.

### Somatotopic organization of higher-order somato-motor cortex in newborns

Beyond the central sulcus, cortical responses to tactile stimulation of the face, hand, and foot were observed throughout motor, pre-motor, somatosensory, parietal, and insular cortex (Supplementary Figure 3). Several whole-body topographic maps were identified in newborn monkeys as early as 11 days of age (Figure 2; Supplementary Figure 4). Surrounding the central sulcus representation (#1), additional topographic representations of the body were identified in both newborns and juveniles (Figure 2). A second gradient of body part representations (#2) was identified within the parietal operculum. Starting from the face representation within the central sulcus, representations of the hand and foot were identified posterior in a smooth gradient. Ventral to this gradient, a third body map (#3) was identified within the insula. Starting from a representation of the face ventral to the central sulcus, hand and foot representations were identified posteriorly. Several additional partial body maps were identified. Anterior to the face representation within the central sulcus, a representation of the hand (#4) was identified within ventral portions of the arcuate sulcus. Regions preferring feet more than hands and the face were not identified in most monkeys within the arcuate. Several representations of the hand and foot (#5-7) were identified adjacent to each other near the midline of cortex. In most monkeys, regions preferring faces more than hands and feet were not identified in midline regions of cortex. The extent of these body representations were similar between newborns and juveniles demonstrating that the large-scale somatotopic representations of the body are already established in week-old monkeys.

**Figure 2.**
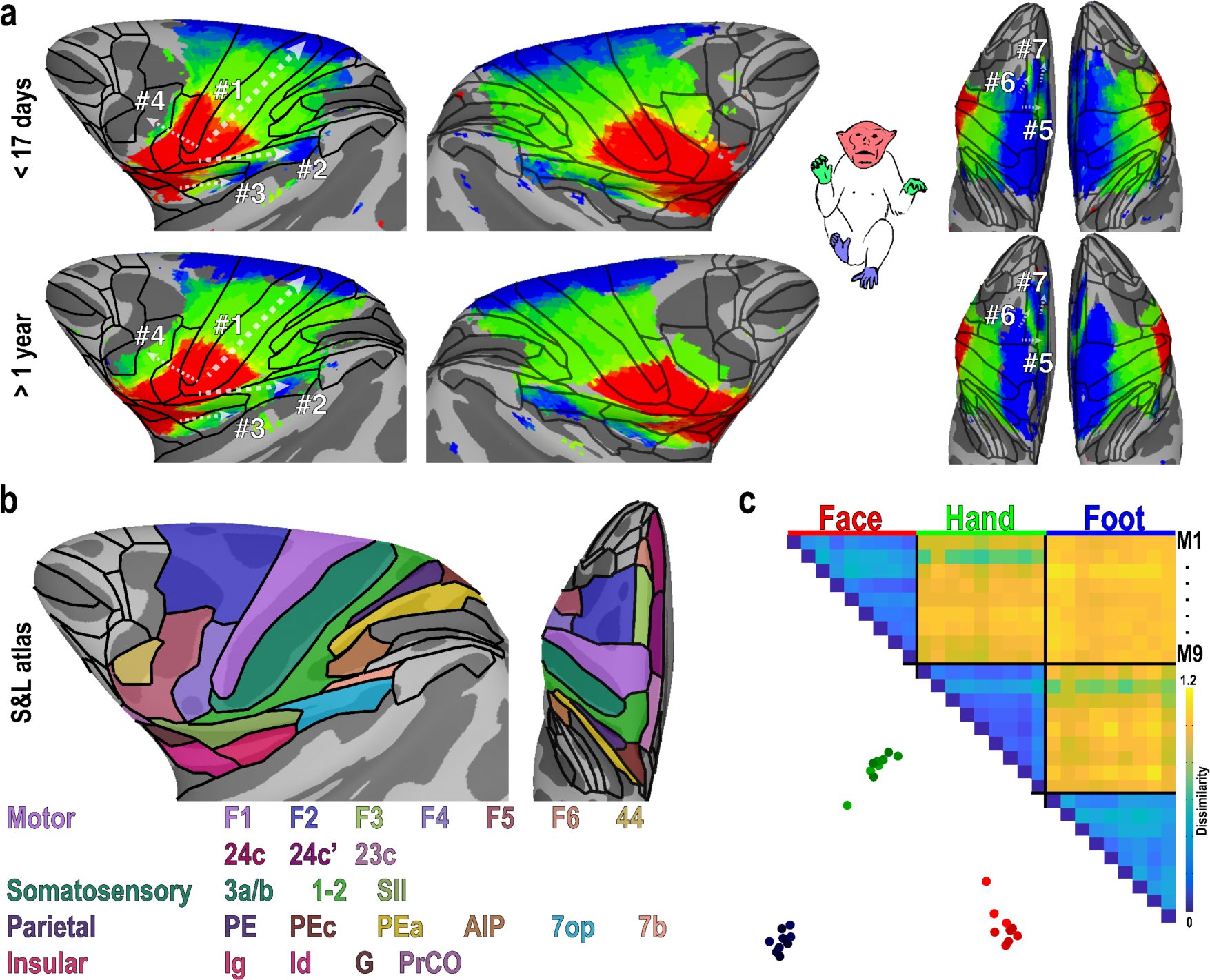
Multiple body maps in newborns and juveniles. (a) Group average body maps of newborn (M1 & M2) and juvenile (M7 & M8) monkeys. Contralateral representations of the face, hand, and foot were found throughout frontal, anterior parietal, and insular cortex in monkeys younger than 17 days old (M1 & M2) and in juveniles older than 1 year (M7 & M8). Seven topographic gradients of body representations were observed in all monkeys. Data threshold at a group average z-statistic > 4 (p < 0.0001, uncorrected) for the strongest IC in each vertex. (b) Somatotopic representations overlapped with 23 areas of the Saleem and Logothetis atlas spanning motor, somatosensory, parietal, and insular cortex. (c) The spatial pattern of face, hand, and foot representations was consistent across all 9 monkeys tested. The dissimilarity matrix (1-corr) and MDS demonstrated that the spatial maps clustered based on body part stimulation, and there was no clear effect of age. For MDS, monkeys were color coded by age from youngest (light) to oldest (dark). Face, hand, and foot representations were color coded red, green, and blue.

**Figure 3.**
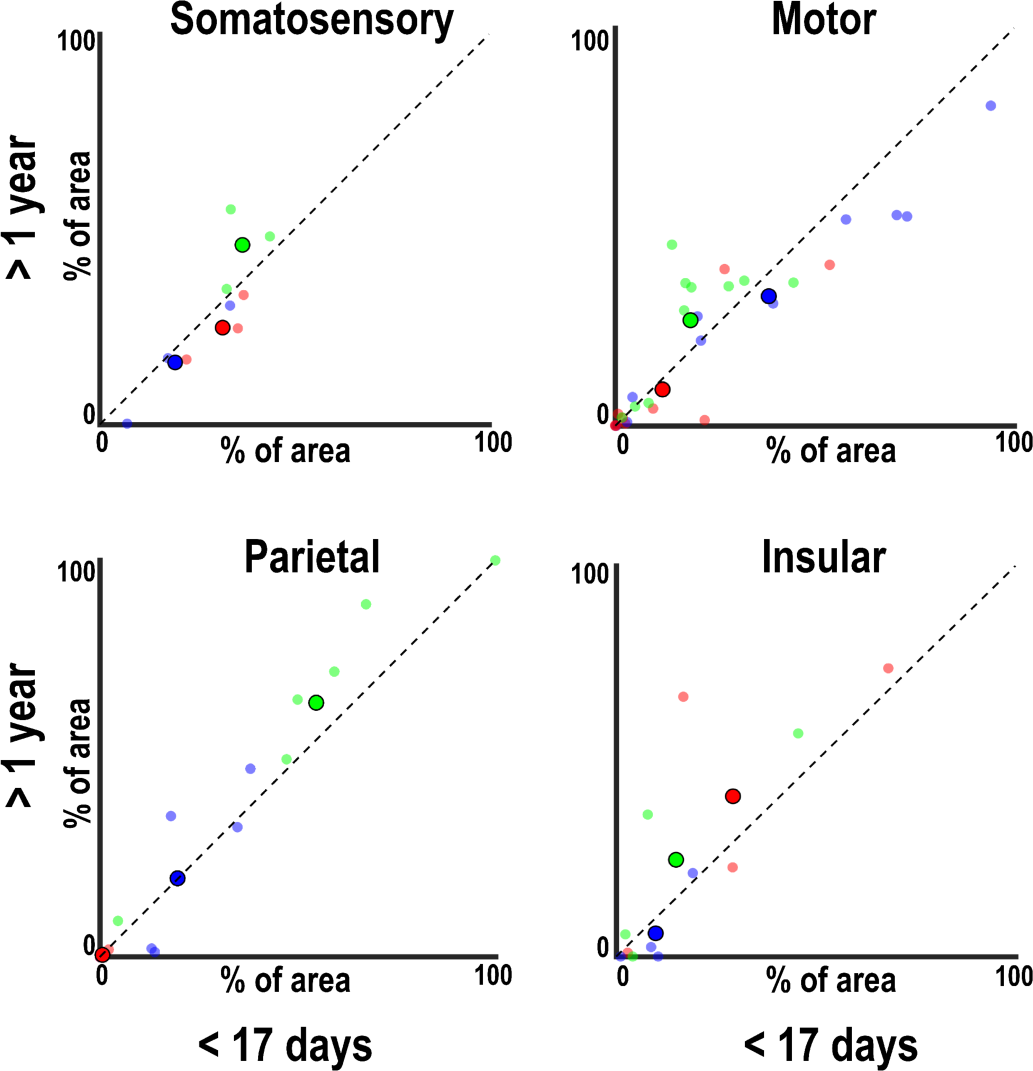
Distribution of body representations across cortex in newborns (M1 & M2) and juveniles (M7 & M8). The cortical area comprising face, hand, and foot representations was comparable between newborns and juveniles for 23 cortical areas identified from the Saleem and Logothetis atlas. Large, black-outlined circles denote average body part representations across somatosensory, motor, parietal, and insular cortical regions. Smaller circles denote average body part representations within individual cortical areas in each cortical region.

**Figure 4.**
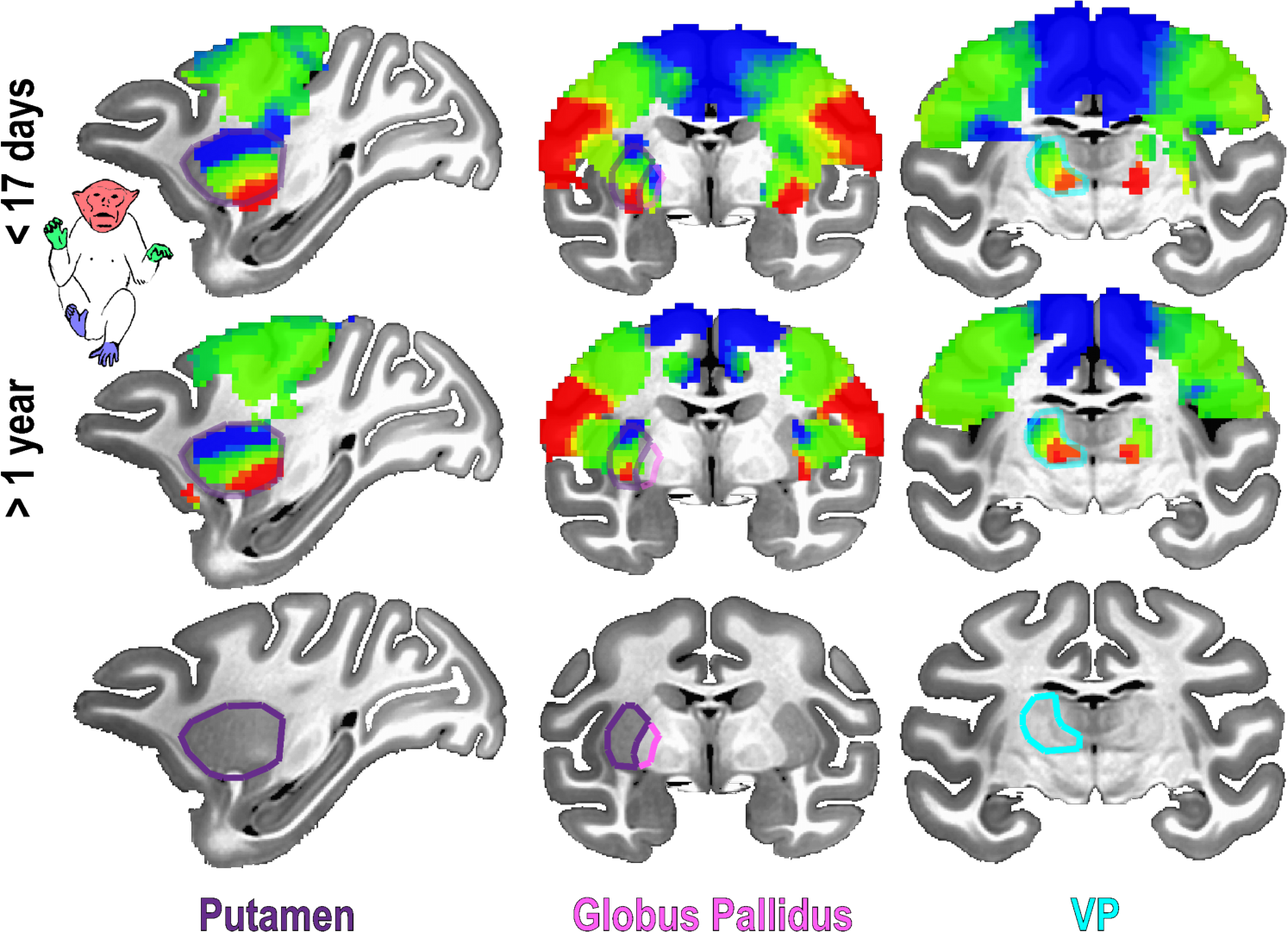
Subcortical somatotopic organization in newborns and juveniles. Somatatopic maps of the contralateral body were found in the putamen, globus pallidus, and ventral posterior nucleus of the thalamus in newborns (M1 & M2), and these maps were comparable to the organization in juvenile monkeys (M7 & M8). Data threshold at a group average z-statistic > 4 (p < 0.0001, uncorrected) for the strongest IC in each vertex.

Topographic gradients of body part representations encompassed several motor, somatosensory, parietal, and insular cortical areas based on an anatomical atlas (Figure 2b). Analogous to the multiple visual-field maps that share a single eccentricity gradient, some of these large-scale topographic gradients may correspond to multiple, distinct body maps that run in parallel across the cortical surface. For example, the topographic gradient within the central sulcus (#1) spanned both primary motor cortex (F1/M1) and primary somatosensory cortex (3a/b), which contain distinct maps of the body ^8, 14^. This topographic gradient also extended into neighboring cortical areas in frontal cortex (F2) and anterior parietal cortex (PE, PEa, Pec, and AIP), which are also known to contain somatotopic representations ^16-20^. Other topographic gradients spanned several architectonic regions. For example, the topographic gradient within and around ventral portions of the arcuate (#4) spanned areas F4, F5 and 44, and the topographic gradient posterior to the central sulcus (#2) spanned SII and 7op, consistent with prior electrophysiology studies in monkeys ^21-24^. Our data suggest a loose correspondence between somatotopic functional maps and anatomy that does not strictly adhere to architectonic borders.

The distribution of body part representations across cortex was consistent across individuals and ages (Figures 2 and 3). As shown in the similarity matrix (Figure 2c), the spatial pattern of ICA maps was more similar across individuals for matched (e.g., face IC of M1 and face IC of M6) than for nonmatched body parts (e.g., face IC of M1 and hand IC of M6). As shown in the MDS plot, data from individual monkeys clustered based on body part representation and did not differentiate between ages (Figure 2c). Further, the distribution of face, hand, and foot representations within cytoarchitectonic regions was similar between newborn and juvenile monkeys for somatosensory, motor, parietal, and insular cortices (Figure 3). These data confirm that stereotypical features of the adult body-map organization are already present in newborns, including overrepresentations of the hand and face in primary somatosensory cortex: a larger portion of cortical real estate in newborns represents the hand and face as compared to the feet in S1 (50% and 34% vs. 14%, respectively). Together, these data suggest that the cortical organization of body maps is already established at birth and is comparable to older monkeys.

### Somatotopic organization of subcortex in newborns

Body part gradient maps were also found throughout subcortex in newborns in the putamen, globus pallidus, area VP, and the anterior pulvinar (Figure 4). Similar to primary somato-motor cortex, each subcortical area contained an inverted topographic representation of the body with the face represented ventrally, hands represented in mid sections and feet dorsally. For thalamic area VP, body representations varied also along the medial-lateral axis with face representations located in ventromedial-most portions and foot representations located in dorsolateral-most portions. Body representations within the anterior pulvinar abutted the body map in area VP and were situated anterior to retinotopic maps within the ventral pulvinar ^25^. The organization of these maps was consistent with prior electrophysiological recordings and tracer studies in monkeys ^26-33^. The distribution of face, hand, and foot representations within each region was similar between newborn and juvenile monkeys. Together, these data demonstrate that somatotopic body maps are already present in subcortical nuclei of newborns.

### Postnatal map refinement

While the large-scale body-map organization of the body is present at birth, the extent of the body represented by each neuron, i.e. its receptive field, may be refined over the course of early development. To track receptive field refinement across early development, we mapped individual digit representations within the cortical hand representation. To maximize the repetitions of digit stimulation and thereby our chance of finding digit representations, we stimulated only the right-hand digits during a full scan session for newborn monkeys (M1, M2). Digits in both hands were mapped in a full scan session for monkeys M3 and M4 and only in the right-hand digits were mapped in M8 at the end of the body mapping scan session. In contrast to the adult-like hand representations found in these newborn monkeys (Figure 2; Supplementary Figures 1 & 2), representations of individual digits were not clearly differentiated. In monkeys younger than 18 days, only small, focal regions representing a few contralateral digits were found within the hand representation of primary somatosensory (3a/b) cortex (Figure 5, left). In monkey M1, representations of the contralateral thumb (red) and index finger (yellow) were found within the left hemisphere. In monkey M2, a representation of the middle finger and weak representations of the thumb and pinkie were found within the left hemisphere. By a few months of age, contralateral representations of each finger could be clearly identified in both primary (3a/b) and secondary (SII) cortex (Figure 5, middle; Supplementary Figure 5). Though evoked responses in these monkeys were relatively weak (< 1% signal change) and similar to newborns (Supplementary Figure 6), the digit representations in 2-7 month old monkeys were more dissimilar than in monkeys younger than 18 days (Figure 5, bottom; Mean Euclidean distance on group average maps = 1.29 (0.04) vs. 0.39 (0.02); paired t-test, t(9) = −10.6449, *p* < 0.0001). By ~2 years, these representations were more robust (Figure 5, right) and the evoked responses were larger than in younger monkeys (Supplementary Figure 6). Interestingly, the digit representations were not any more differentiated than in the 2-7 month old monkeys (Figure 5, bottom; Mean Euclidean distance on group average maps = 1.28 (0.05); paired t-test, t(9) = 0.22; *p* = 0.83). Together, these data indicate that the receptive fields of hand digits are coarse at birth, and fine-grained spatial precision is refined over the first 2 years.

**Figure 5.**
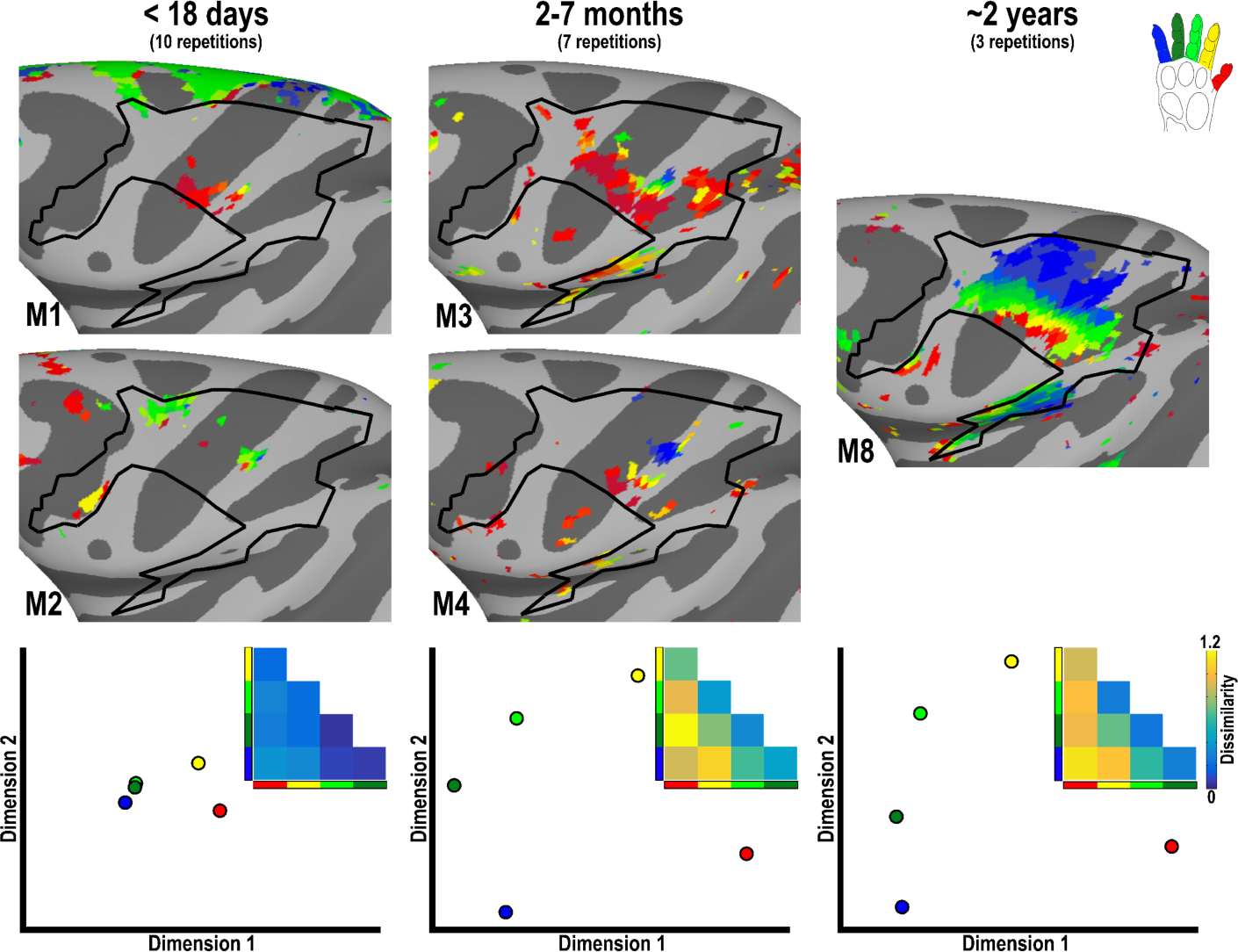
Digit representations in newborns and juveniles. With the exception of the thumb and index fingers, monkeys less than 18 days old (M1 & M2) lacked spatially distinct representations of contralateral fingers in somato-motor cortex. In monkeys 2-7 months old (M3 & M4), representations of each digit were identified in primary (3a/b) and secondary (SII) somatosensory cortex. At ~2 years-old (M8), digit representations appeared more robust in primary and secondary somatosensory cortex and were also found within motor cortex. Data threshold of p < 0.0001, uncorrected.

**Figure 6.**
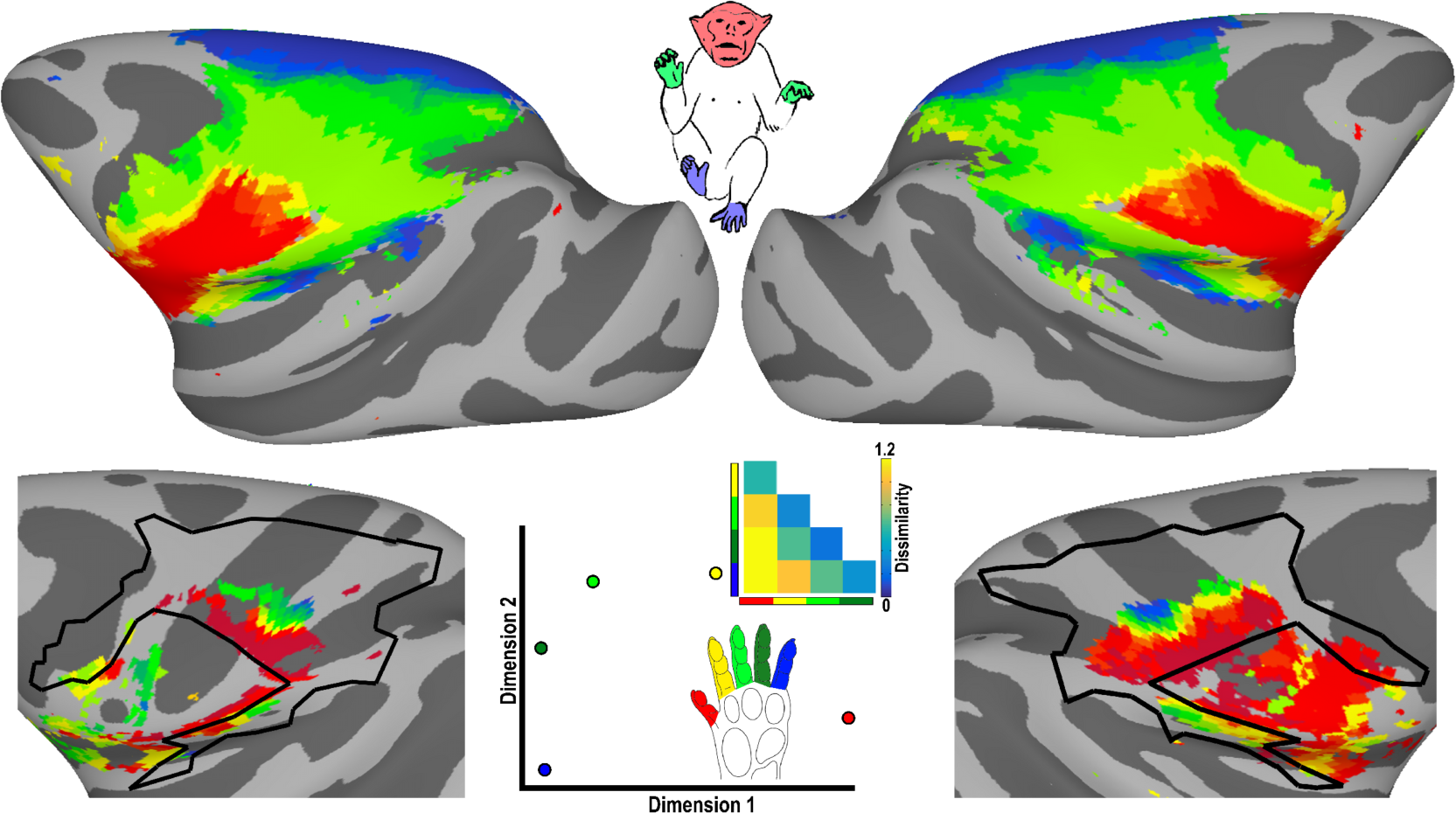
No clear somatotopic reorganization in monkeys raised under visual form deprivation. Somatotopic maps were found in two monkeys (M5 & M6) raised without visual form experience for the first year of life. These monkeys primarily rely on touch for navigation and interacting with their environment. Despite this massive shift in sensory experience relative to control monkeys, the somatotopic organization was indistinguishable from controls. Somatosensory responses were not observed in occipital cortex, suggesting that such drastic changes in early sensory experience were not sufficient to cause large-scale cross-modal re-wiring. Body and digit mapping data threshold of p < 0.0001, uncorrected.

### Lack of cross-modal remapping in sensory deprivation

Despite the presence of an adult-like large-scale somatotopic organization at birth, the early topographic organization is nevertheless more plastic than in adults, being capable of dramatic alteration following injury to peripheral input ^34-37^ or through dramatic changes in early somato-motor experience ^38^. The extent to which somatotopic representations are modifiable from alterations of other sensory modalities remains to be resolved. Studies in congenitally blind humans have shown somatosensory-evoked responses in visual cortex ^39^, suggesting that early deprivation in one sensory modality can lead to re-wiring, such that other sensory modalities take over cortical territory typically performing computations related to the deprived modality. Here, we asked whether such cross-modal re-wiring can be seen in monkeys and whether early experience (or lack thereof) is sufficient for driving such large-scale changes. As part of a separate experiment, we raised two monkeys for the first year of life with visual experience restricted to diffuse light. Without form vision, behaviorally, touch became the dominant sensory modality in these animals. In contrast to control monkeys that rely on vision for navigation and identifying objects in their environment, these monkeys predominantly relied on touch both during and after the deprivation period. For example, when placed in an unfamiliar environment, normally reared monkeys will visually scan the layout. In contrast, these two monkeys surveyed the layout by making several passes around the perimeter, feeling the walls and floor with their hands. We asked whether there was a corresponding shift in the neural architecture dedicated to processing touch. In contrast to the stark behavioral shift, we found that somatotopic representations in these monkeys were comparable to age-matched control monkeys (Figure 6, top). Somatotopic representations were found throughout the central sulcus, frontal and parietal cortices as well as subcortex. Somatosensory-evoked responses were not found in either occipital or temporal cortex that typically responds to visual input. The extent of receptive fields also appeared comparable to age-matched control monkeys. Individual digit representations were identified in visually deprived monkeys (Figure 6, bottom). The topographic organization of the hand digits was comparable to controls of similar ages. Digit representations in visually deprived monkeys were not significantly more (or less) dissimilar than 2-7 month old monkeys (Mean Euclidean distance on group average maps = 1.29 (0.05); paired t-test, t(9) = 0.07; *p* = 0.94). Together, these data suggest that an abnormal prioritization of touch starting at birth is insufficient to generate a large-scale somatotopic reorganization across sensory systems.

## Discussion

Somatosensory and motor systems were found to be functionally organized to a remarkable degree in awake newborn macaques as young as 11 days postnatal. Here, we found spatially distinct activity in response to tactile stimulation of the contralateral face, hand, and foot within primary somato-motor cortex. The organization of these body maps was comparable to that of older monkeys. A topographic gradient of body representations spanned the dorsal-ventral extent of the central sulcus from a representation of the face at the ventral tip to an adjacent hand representation and to a representation of the foot at the dorsal tip. This gradient covered somatotopic areas 3a/b and 1-2 as well as primary motor area M1/F1 and is consistent with prior microelectrode recordings within the primary somatosensory ^7-9^ and motor cortex ^2, 13, 14^ of adults monkeys. Compared with foot representations, the face and hand representations covered a greater extent of the cortical surface, consistent with the over-representation classically illustrated in homunculi. The presence of somatotopic organization in newborns may appear in contrast to prior electrophysiological studies that found neurons within primary somatosensory cortex of newborn monkeys to be unresponsive to tactile stimulation ^5^. However, these prior recordings were done under anesthesia, which may have dampened evoked responses. Interestingly, the magnitude of fMRI evoked responses to stimulation of the face, hand and foot was weaker in our younger monkeys (up to 146 days) than in older monkeys (Supplementary Figure 1), suggesting that while the somatotopic organization is adult-like, response properties remain immature for the first several months. Our data are consistent with a recent paper that showed spatially distinct representations of the face, hand, and ankle within primary somato-motor cortex in preterm humans ^40^. Our data demonstrate that the large-scale topographic organization of primary somato-motor cortex is adult like at birth and indicates that postnatal experience is not needed for their formation. These maps likely form from a combination of molecular cues and activity-dependent sorting ^41^, though in utero tactile and motor experience could also play a role.

Beyond primary somato-motor cortex, spatially distinct representations of the contralateral face, hand, and foot were observed throughout frontal, anterior parietal, and insular cortex as well as in subcortex. In total, seven body-part gradients were identified across the cortical surface and within subcortical structures. At least some of these gradients likely comprise multiple body maps that run parallel to each other, and thereby were indistinguishable in the current data. The organization of these somatotopic maps are in good agreement with prior findings from anatomical tracer and electrophysiological studies ^16-24, 26-33^. To our knowledge, our data constitute the first comprehensive account of somatotopic organization across the entire macaque brain. Despite the potential for considerable postnatal re-organization and refinement, the somatotopic organization of these areas in newborns was indistinguishable from juvenile monkeys, indicating that the large-scale somatotopic organization of the entire brain is already established at birth.

Despite an extensive “adult-like” somatotopic organization, the fine-scale representations of individual hand digits were not well differentiated in newborn monkeys. In comparison, complete maps of hand digits were present in primary and secondary somatosensory cortex in 2-7 month old monkeys. This refinement of the newborn somatotopic organization coincides with the maturation of fine motor skills. At birth, monkeys are behaviorally immature in motor function and lack adult-like precision and coordination. Motor movements become more precise and coordinated over the first several months ^42, 43^. Thus, while the large-scale somatotopic organization of the newborn brain is indistinguishable from adults, refinement of receptive fields occurs over the course of the first year, paralleling the refinement of fine motor movements.

While early somato-motor experience refined the somatotopic organization, the organization of somato-motor systems was unaffected by dramatic shifts in early sensory experience. The somato-motor system is capable of massive reorganization from early somatosensory loss ^34-37, 44^. In humans with early vision loss, visual cortex can respond to other sensory modalities such as touch ^38, 39, 45^. We tested whether such cross-modal plasticity occurred in visually deprived monkeys. In contrast to control monkeys that primarily relied on vision for navigation and interacting with their environment, visually deprived monkeys relied on touch even after the deprivation period. Despite the dominance of touch over other sensor modalities, the large-scale somatotopic organization in these monkeys was indistinguishable from that in control monkeys, demonstrating that early sensory deprivation alone was insufficient to induce large-scale cross-modal reorganization. In contrast to prior findings in early-blind humans ^38, 39, 45^, in whom tactile stimulation evoked activity within occipital cortex, no increased activity in occipital or temporal cortex was observed in these monkeys from face, foot, or hand stimulation. The large shift in these monkeys’ sensory experience was not driven by trauma to the eye or peripheral pathways that typically induce sensory loss in human studies, suggesting that spontaneous activity within intact anatomical pathways (or simply the experience of diffuse light) may limit the degree of cross-modal plasticity. Thus, the presence of topographic maps at birth provides constraints on plasticity that likely serve an important role in establishing and maintaining sensory representations throughout development.

A proto-architecture of somatotopy may support subsequent experience-driven refinement throughout the somato-motor system. In addition to representations of individual body parts, mature motor cortex contains functional domains specialized for the execution of coordinated movements across multiple joints. These domains emphasize different complex, ethologically relevant, categories of action ^15^ and are localized to stereotypical locations of somatotopic maps across individuals (Figure 7). For example, a domain that supports self-feeding (i.e. bringing hand to mouth) is localized to ventral regions of motor cortex spanning representations of the face and hands and another domain that supports climbing and leaping is localized to dorsal regions of motor cortex spanning representations of the feet and hands. Though the developmental timeline of these domains has yet to be probed, such functional specialization can emerge in self-organizing simulated networks over the course of training on a set of complex movements, without requiring domain-specific priors ^46^. We speculate that the early somatotopic organization observed in newborn monkeys provides the scaffolding for the subsequent development of these functional domains. This organization parallels our recent findings that the entire visual system is retinotopically organized at birth ^47^ and likely supports experience-driven development of behaviorally relevant domains, such as those that selectively respond to text or faces ^48^. Together, these findings illustrate that topographic maps of sensory space are a fundamental and pervasive (Figure 8) organizing principle of the brain, and they play an important role in development by guiding and constraining experience-driven refinement and specialization.

**Figure 7.**
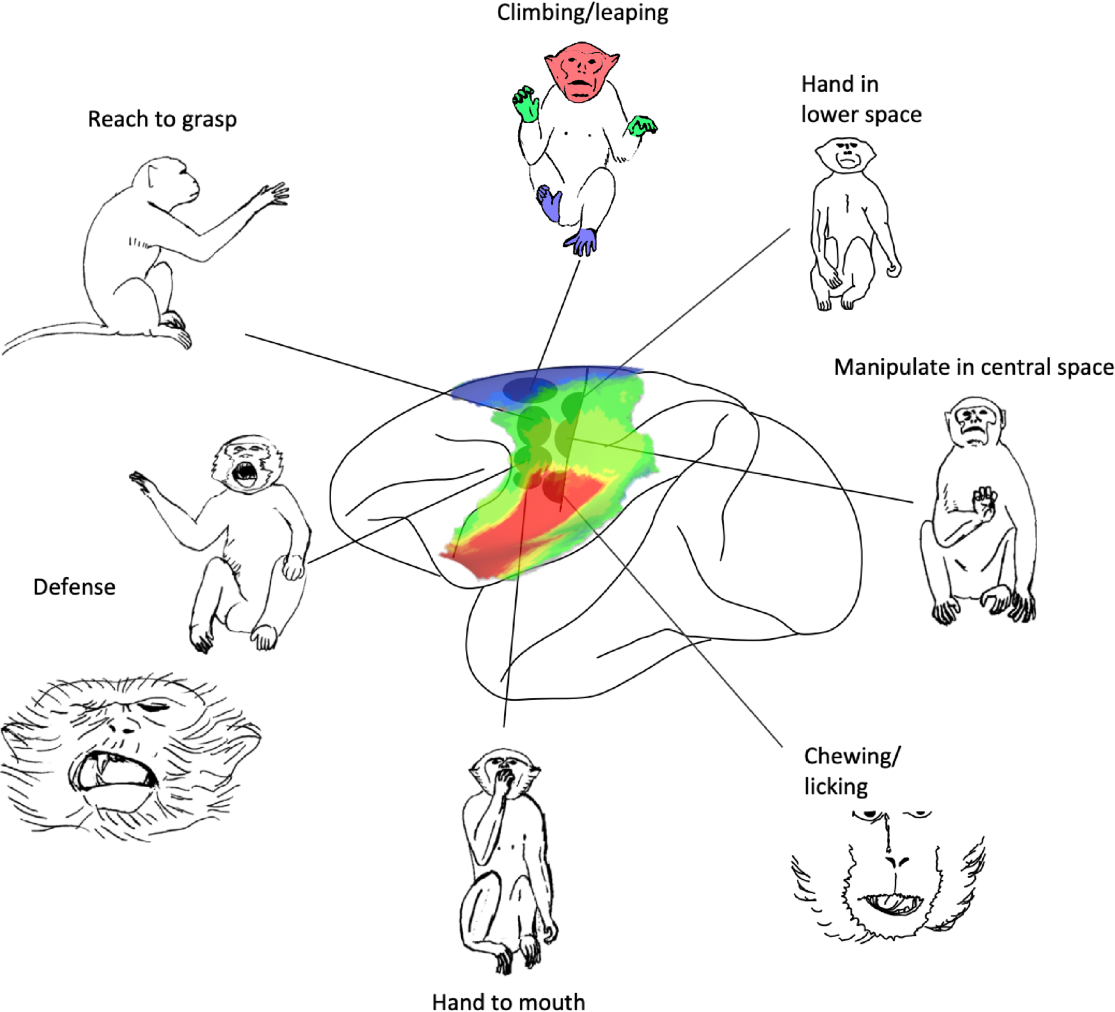
Schematic of the relationship between somatotopy and action domains. Image adapted from Graziano 2016 ^15^. Somatotopic maps were manually aligned to this image based on the anatomical landmarks of the central, arcuate, and insular sulci.

**Figure 8.**
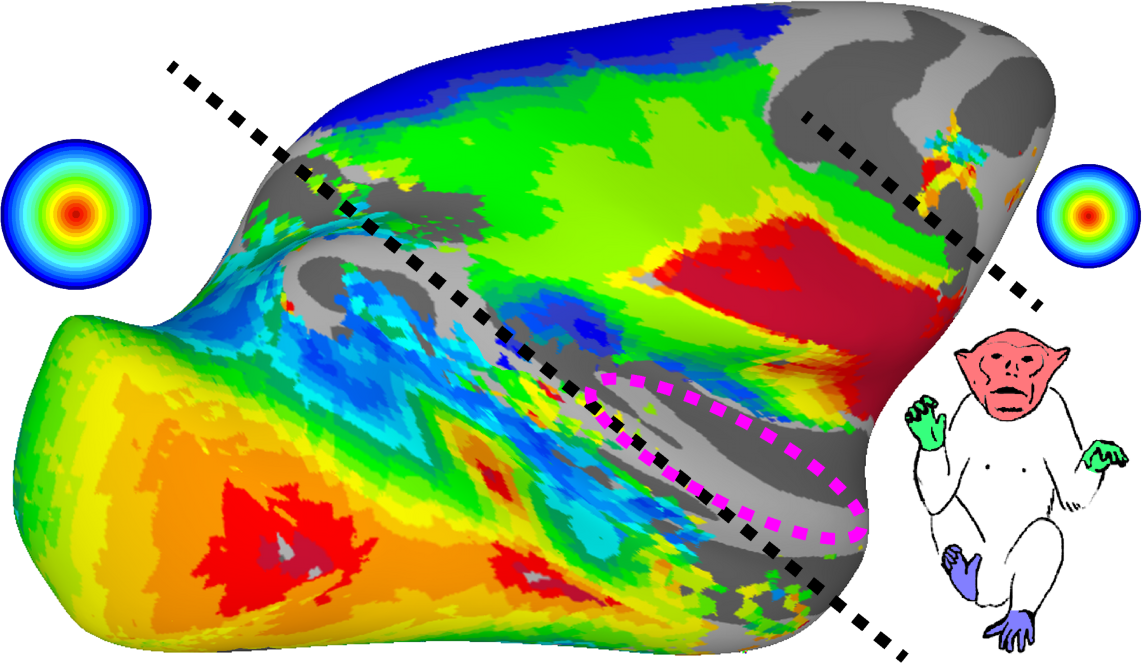
Topographic organization across the cortical surface. Topographic representations of our environment cover most of the cortical surface. As illustrated by a group average (n = 6) eccentricity map ^25^, the posterior half of the brain and an anterior part of the arcuate sulcus (FEF) comprise retinotopic representations of visual space. As illustrated by a group average (n = 8) body map, anterior parietal and frontal cortex comprise somatotopic representations of the body. These retinotopic ^47^ and somatotopic maps (current study) are present at birth. Black dashed lines differentiate retinotopic and somatotopic maps. Retinotopic and somatotopic representations partially overlap within anterior parietal cortex. As illustrated by the magenta dashed circle, several tonotopic maps span the lower bank of the lateral sulcus ^49^.

## MATERIALS AND METHODS

### Monkeys

Functional MRI studies were carried out on 9 Macaca mulattas, 4 female and 5 male, between the ages of 11 days and 981 days. All procedures were approved by the Harvard Medical School Animal Care and Use Committee and conformed to National Institutes of Health guidelines for the humane care and use of laboratory animals. All monkeys were born in our laboratory. One monkey (M9) was co-housed with their mother in a room with other monkeys for the first 4 months, then co-housed with other juveniles, also in a room with other monkeys. As part of separate experiments, all other monkeys were hand reared by humans for the first year, then were co-housed with other juveniles. Two of the hand-reared monkeys (M5 & M6) were raised under conditions of visual form deprivation via eye lid suturing for the first year. Experiments were conducted in these two monkeys after eyelid re-opening. For scanning all monkeys were alert, and their heads were immobilized using a foam-padded helmet with a chinstrap that delivered juice. The monkeys were scanned in a primate chair that allowed them to move their bodies and limbs freely, but their heads were restrained in a forward-looking position by the padded helmet. The monkeys were rewarded with juice during scanning.

### Somatotopic Mapping

For monkeys M1 – M8, somatotopic mapping was performed using air puffs for face stimulation and gentle manual stroking for body-part stimulation. For monkey M9, hand and foot responses were measured via air puffs. This method was found to be non-optimal because the tubes that delivered air puffs kinked during the scan due to monkey movement resulting in unreliable stimulation. Body maps are shown for monkey M9, but because of this different method of body part stimulation, data were not included in the direct comparisons between newborns and juvenile. For the two youngest monkeys (M1 & M2), hand, foot, and face responses were measured for the right side of the body only, to increase the number of repetitions collected. For monkeys M3 – M8, hand, foot, lower back, and face responses were measured bilaterally. Each scan comprised blocks of stimulation of individual body parts; block length was 20 seconds, with 20 seconds of no-stimulation interleaved. Seven monkeys (M1 – M6, M8) participated in a finger mapping experiment where responses to manual stroking of the thumb, index, middle, ring, and pinkie were measured. For the two youngest monkeys (M1 & M2), responses were measured only to the right hand, again to increase the number of repetitions collected. For the other five monkeys (M3 – M6, M8), responses to both hands were mapped. Monkeys were rewarded with Juice every couple of seconds throughout the scan.

### Scanning

Monkeys were scanned in a 3-T TimTrio scanner with an AC88 gradient insert using 4-channel surface coils (custom made by Azma Maryam at the Martinos Imaging Center). Each somatotopic scan session consisted of 10-12 functional scans. We used a repetition time (TR) of 2 seconds, echo time (TE) of 13ms, flip angle of 72ᵒ, iPAT = 2, 1mm isotropic voxels, matrix size 96×96mm, 67 contiguous sagittal slices. To enhance contrast^50, 51^, we injected 12 mg/kg monocrystalline iron oxide nanoparticles (Feraheme, AMAG Parmaceuticals, Cambridge, MA) in the saphenous vein just before scanning.

### General preprocessing

Functional scan data were analyzed using Analysis of Functional NeuroImages (AFNI; RRID:nif-0000-00259)^52^, SUMA^53^, Freesurfer (Freesurfer; RRID:nif-0000-00304)^54, 55^, JIP Analysis Toolkit (written by Joseph Mandeville), and MATLAB (Mathworks, RRID:nlx_153890). Each scan session for each monkey was analyzed separately. All images from each scan session were AFNI-aligned to a single timepoint for that session, detrended and motion corrected. Data were spatially filtered using a Gaussian filter of 2 mm full-width at half-maximum (FWHM) to increase the signal-to-noise ratio (SNR) while preserving spatial specificity. Each scan was normalized to its mean. Data were registered using a two-step linear then non-linear alignment approach (JIP analysis toolkit) to a standard anatomical template NMT; ^56^ for all monkeys. First, a 12-parameter linear registration was performed between the mean EPI image for a given session and a high-resolution anatomical image. Next, a nonlinear, diffeomorphic registration was conducted. To improve registration accuracy of ventral cortex, masks were manually drawn that excluded the cerebellum for both EPIs and anatomical images prior to registration.

### Regression Analysis

A multiple regression analysis (AFNI’s 3dDeconvolve ^52^) in the framework of a general linear model ^57^ was performed on the somatotopic mapping experiments for each monkey separately. Each stimulus condition was modeled with a MION-based hemodynamic response function ^50^. Additional regressors that accounted for variance due to baseline shifts between time series, linear drifts, and head motion parameter estimates were also included in the regression model. Due to the time-course normalization, beta coefficients were scaled to reflect percent signal change. Since MION inverts the signal, the sign of beta values were inverted to follow normal fMRI conventions of increased activity represented by positive values. Maps of beta coefficients for each body part stimulated set to a threshold of *p* < 0.0001 (uncorrected).

### Independent Component Analysis (ICA)

A 2D (space × time) ICA (FSL’s MELODIC) was performed on the concatenated runs for each scan session. ICA is a special case of blind source separation where signals throughout the brain are parsed into statistically independent components. This analysis decomposes each space × time matrix into pairs of temporal and spatial subcomponents. This analysis assumes that the subcomponents are non-Gaussian signals and are statistically independent from each other. The analysis was restricted to voxels that fell within a whole brain mask. Between 15-40 components were identified in each scan session. For each scan session, subcomponent spatial maps that were focal to the face, hand, and foot regions of somatosensory cortex were identified in each monkey. These spatial maps were threshold at z > 4.00 (*p* < 0.0001, uncorrected).

### Body Map Analysis

Topographic maps of the body were derived from the face, hand, and foot spatial IC maps. For each voxel, the z-statistics of the face, hand, and foot ICAs were extracted and were normalized to the maximal response. These normalized responses were linearly interpolated into a 100-point 3-dimensional scale. The first, second, and third dimensions corresponded to the weighting of face, hand, and foot responses, respectively. e.g., ‘1 0 0’ corresponded to a weighting of 100% face response and 0% foot and hand. ‘0 1 0’ corresponded to a weighting of 100% hand response and 0% face and foot. Voxels were threshold such that the z-statistic of the peak voxel > 4.00 (*p*< 0.0001, uncorrected).

### Topographic similarity analyses

The spatial similarity of face, hand, and foot representations was compared by (Pearson) correlating the IC maps across monkeys then averaging across hemispheres. This yielded a symmetric 27 × 27 (9 monkeys × 3 body part) similarity matrix (Figure 2c). Classical multidimensional scaling (MDS) was applied on the Euclidean distances between these correlations and the first two principal dimensions were visualized. The spatial similarity of digit representations was compared by correlating the IC maps of thumb, pointed, middle, index, and pinkie fingers across monkeys. Because there was a clear difference in the digit maps between newborns and juveniles, data were grouped into 3 age ranges (< 18 days, 2-7 months, and > 1 year) and MDS was applied on each group separately. For each group, the first two components were visualized (Figure 5, bottom; Figure 6). The distances between digit representations were compared between groups with non-paired, two-tailed t-tests.

### Comparison to atlas

To directly compare visual field maps across monkeys, each monkey’s data was aligned to a standard template (NMT) surface using nonlinear registration (JIP Analysis Toolkit). Group average maps were compared with the borders of the Saleem and Logothesis macaque atlas ^58^ that was aligned to the NMT anatomy ^56^.

## Authors contributions

MSL, PFS and MJA scanned monkeys; MJA analyzed the data; MJA wrote the paper.

## Acknowledgments

This work was supported by NIH grants RO1 EY 25670, P30 EY 12196, and F32 EY 24187. This research was carried out in part at the Athinoula A. Martinos Center for Biomedical Imaging at the Massachusetts General Hospital, using resources provided by the Center for Functional Neuroimaging Technologies, P41EB015896, a P41 Biotechnology Resource Grant supported by the National Institute of Biomedical Imaging and Bioengineering (NIBIB), National Institutes of Health, and NIH Shared Instrumentation Grant S10RR021110.

