## Supplementary Figures for "Body-map proto-organization in newborn macaques"

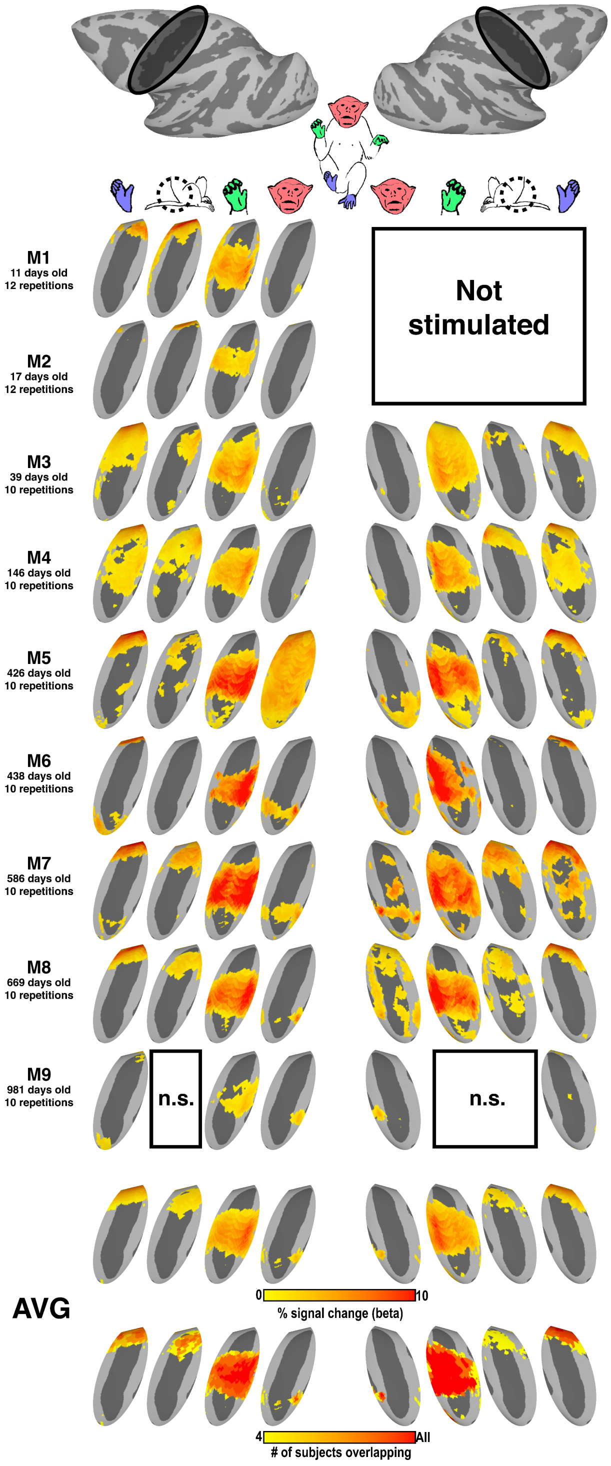


*Supplementary Figure 1. Task-evoked activity in primary somato-motor cortex from stimulation of the contralateral face, hand, lower back, and foot in individual monkeys. Group average evoked activity are shown at the bottom along with a conjunction analysis showing the number of monkeys for which each vertex was significantly activated. Individual monkey data threshold at p < 0.0001, uncorrected. For group average data, only vertices where significant activity was identified in 4 or more monkeys are shown. Only the right side of the body was stimulated for monkeys M1 and M2. The lower back was not stimulated for monkey M9 and a kink in the air tub prevented stimulation of the left hand in monkey M9. See Figure 1 for additional conventions.*

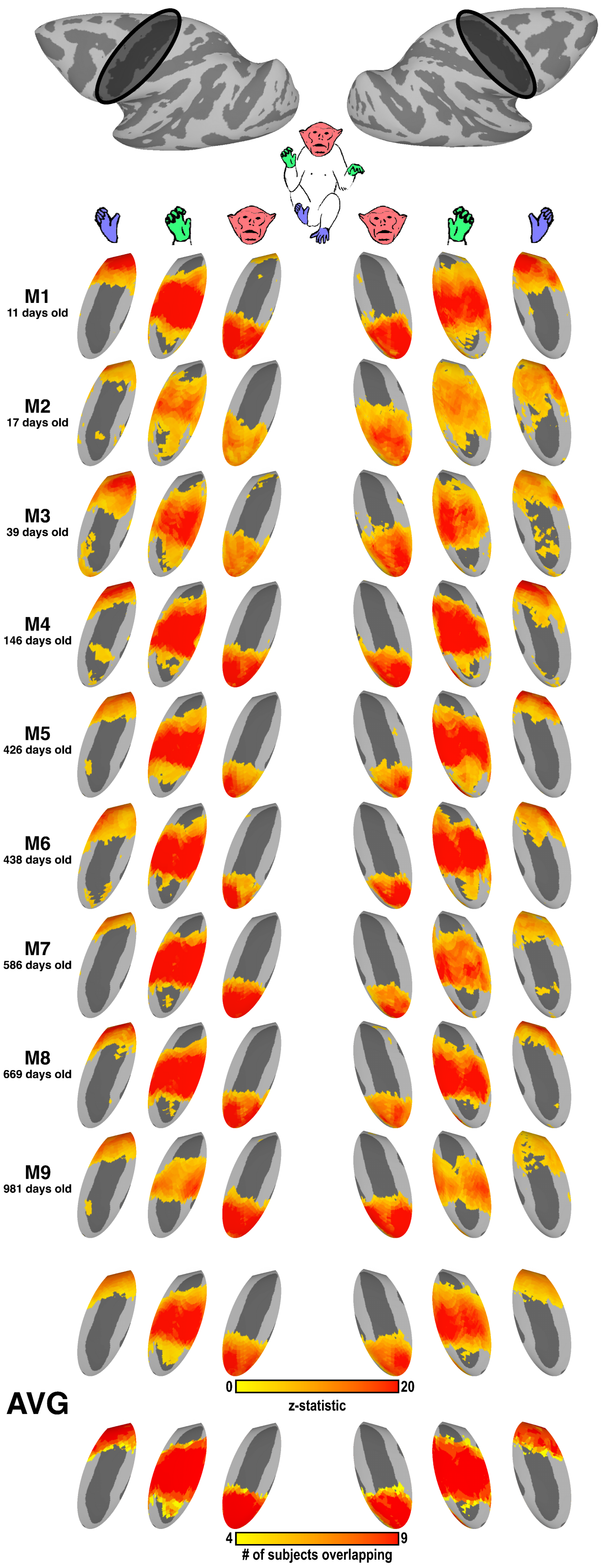


*Supplementary Figure 2. Independent components from stimulation of contralateral body in individual monkeys in primary somato-motor cortex. Independent component maps that correspond to activity within face, hand, and foot representations within the central sulcus are shown. Group average evoked activity and a conjunction analysis are shown at the bottom. Individual ICs threshold at z-statistic > 4 (p < 0.0001, uncorrected). For group average data, only vertices where significant activity was identified in 4 or more monkeys are shown. See Figure 1 and Supplementary Figure 1 for additional conventions.*

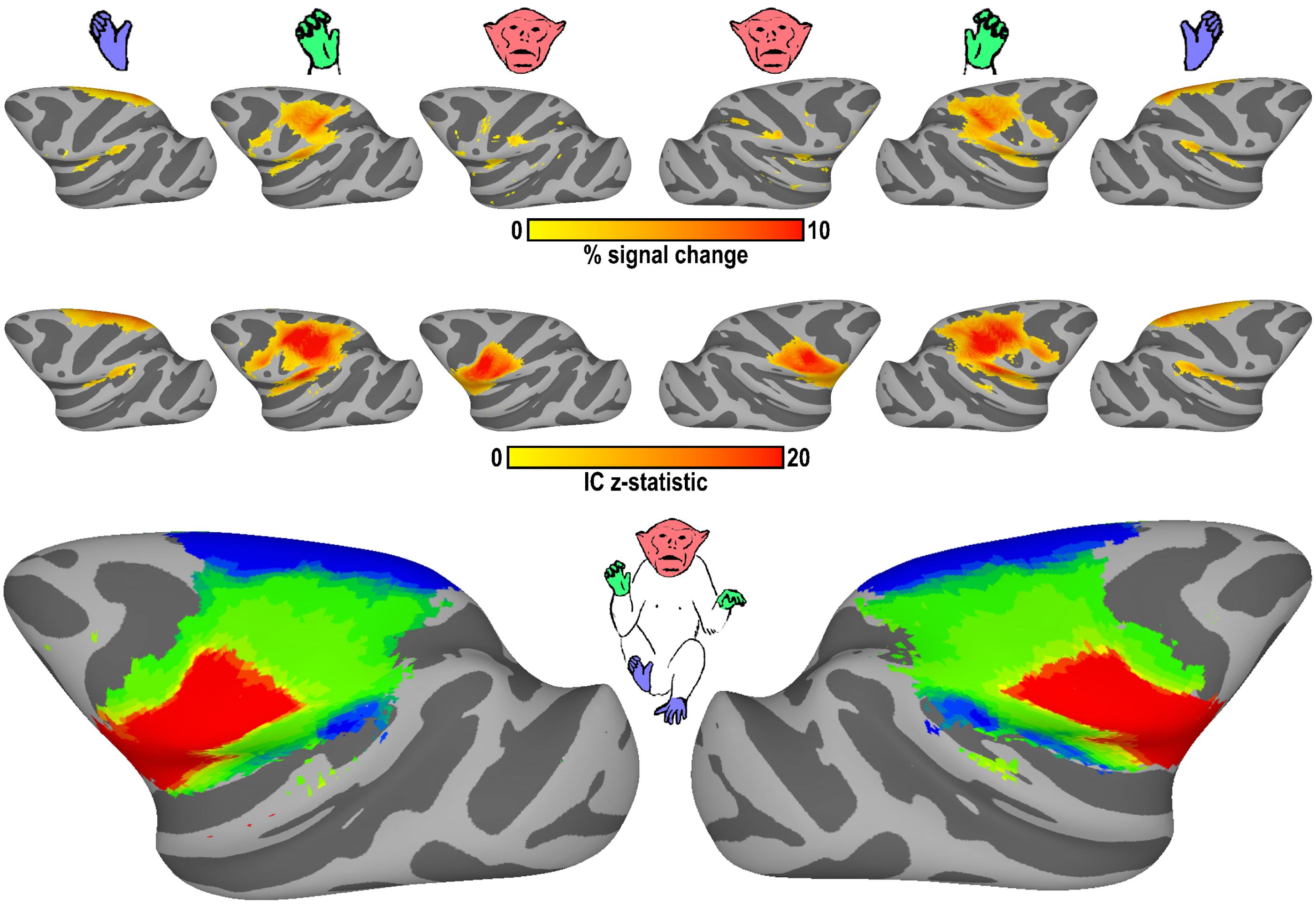


*Supplementary Figure 3. Task-evoked activity and independent components across the cortical surface from stimulation of contralateral body. (top) Group average (n=9 for right hemisphere and n=7 for left hemisphere) task-evoked activity from stimulation of the contralateral hand, foot, and face. (middle) Group average (n=9) independent components corresponding to the face, hand, and foot representations. (bottom) Topographic body map from group-average IC maps. Only vertices where significant activity (z-statistic > 4, p < 0.0001, uncorrected) was identified in 4 or more monkeys are shown. See Figures 1 and 2 for additional conventions.*

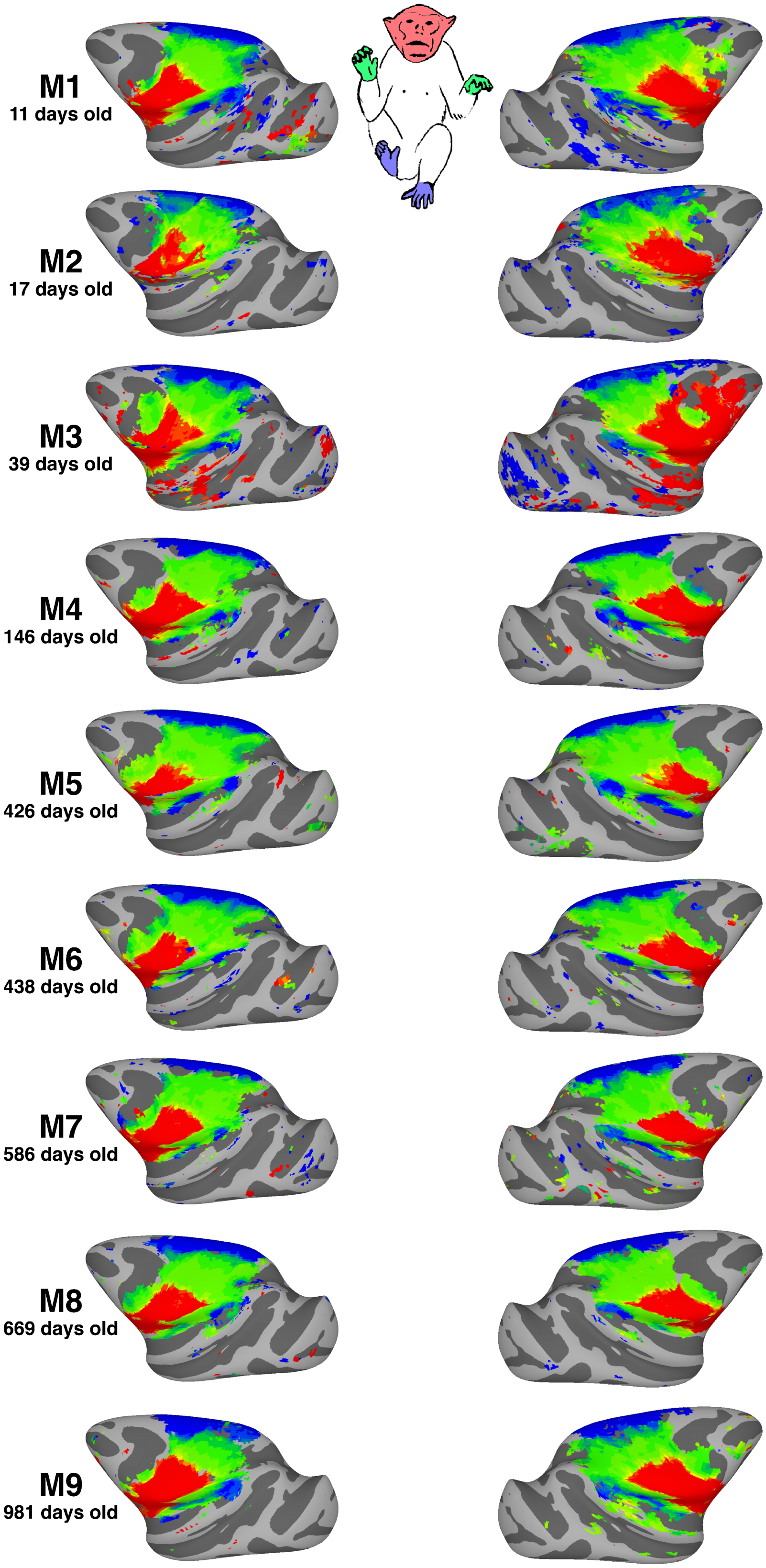


*Supplementary Figure 4. Topographic body map across the cortical surface for each monkey. Topographic gradient maps were calculated from the foot, hand, and face IC maps in each monkey. Data threshold at a z-statistic > 4 (p < 0.0001, uncorrected) for the strongest IC in each vertex. See Figures 1 and 2 for additional conventions.*

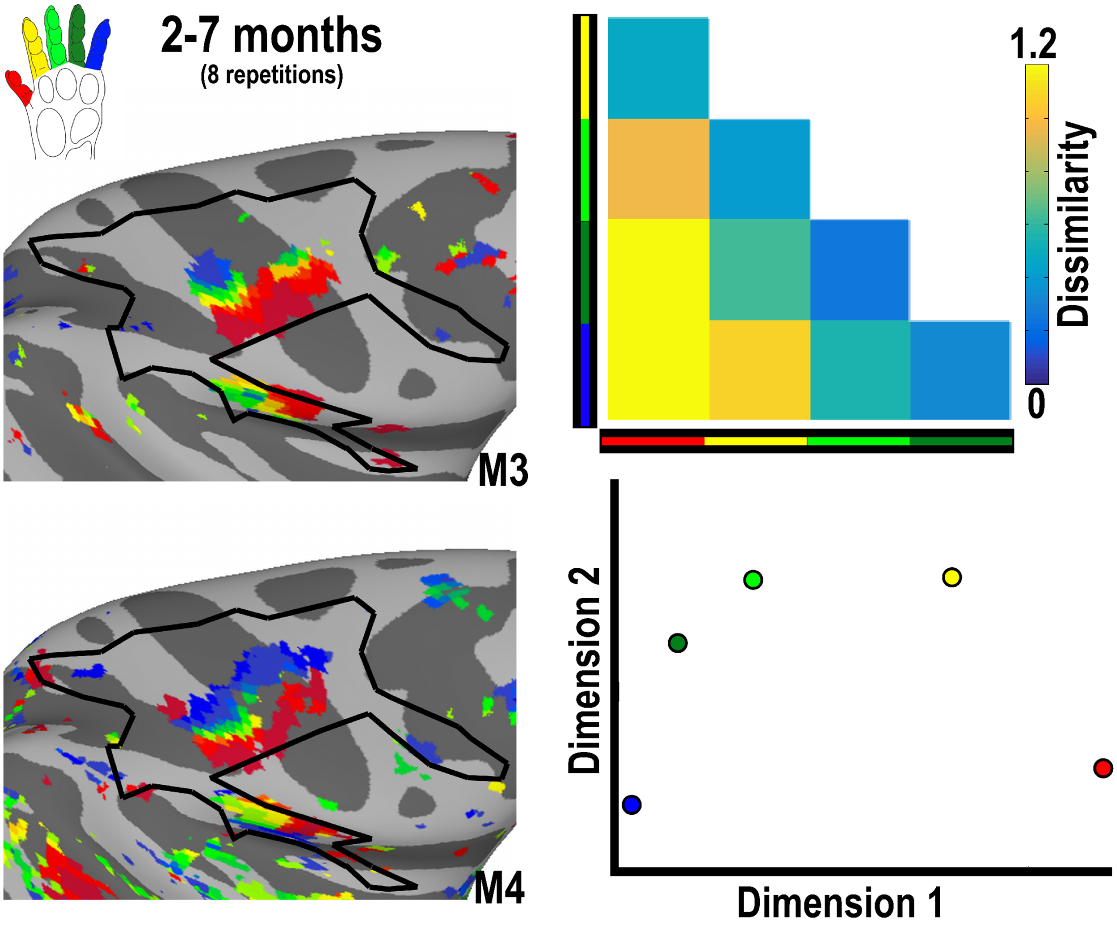


*Supplementary Figure 5. Digit maps in the right hemisphere of 2-7 month old monkeys from contralateral stimulation. Representations of each digit were identified in primary (3a/b) and secondary (SII) somatosensory cortex. Data threshold of p < 0.0001. See Figure 5 for additional conventions.*

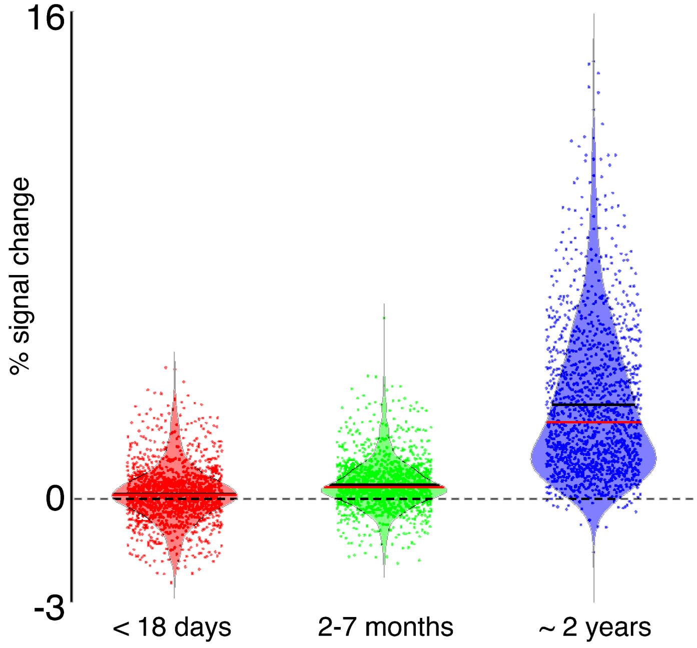


*Supplementary Figure 6. Distribution of evoked responses from digit stimulation in the hand representation primary somato-motor cortex for newborns, 2-7 month old, and ~2-year old monkeys. Red line illustrates median, Black line illustrates mean.*
